# Scc2-mediated loading of cohesin onto chromosomes in G1 yeast cells is insufficient to build cohesion during S phase

**DOI:** 10.1101/123596

**Authors:** Kim A Nasmyth

**Affiliations:** Department of Biochemistry, Oxford University, South Parks Road, Oxford, OX1 3QU, UK

## Abstract

Sister chromatids are held together from their replication until mitosis. Sister chromatid cohesion is mediated by the ring-shaped cohesin complex and it is thought that cohesin holds sister chromatids together by entrapping sister DNAs within the cohesin ring (Haering et al., 2008). However, how this occurs is not well understood. Because cohesin binds to DNA prior to replication it is possible that the replication fork passes through the lumen of the ring thereby placing replicated sisters inside cohesin rings. If this is the case, loading of cohesin in the G1 phase may be sufficient to build cohesion.

We show here that Scc2, a cohesin subunit required for loading cohesin onto chromosomes *de novo*, is necessary for establishment of cohesion even after Scc2-mediated loading has already taken place during late G1 or early S phase. Our results challenge a previous conclusion based on related experiments whereby Scc2 was found not to be required for cohesion establishment during S phase (Lengronne et al., 2006).

## Introduction

The notion that sister chromatid cohesion is mediated by co-entrapment of sister chromatin fibres inside individual tripartite cohesin rings (Haering et al., 2008) raises the question of how such structures are generated during S phase. Are they created *de novo* following passage of replication forks, or from rings that had previously entrapped unreplicated fibres? If the latter were true, then co-entrapment could in principle arise from passage of replication forks through cohesin rings. This would then explain in simple terms why only sister DNAs are co-entrapped. If this is indeed what happens and if as currently believed Scc2 is merely involved in loading cohesin onto chromosomes in the first place, establishment of cohesion during S phase should take place in the absence of Scc2 function - if cohesin had already loaded onto chromatin.

To test this prediction, we used the temperature sensitive *scc2-4* allele to modulate Scc2 activity as *S. cerevisiae* cells undergo S phase.

## Results

When cells containing the thermosensitive (ts) *scc2-4* (K15021) allele, that are arrested in early G1 using α-factor pheromone at the permissive temperature (23°C) are released into medium lacking the pheromone in the absence of APC/C^Cdc20^ activity, cells undergo S phase and subsequently arrest in metaphase with sister chromatids held together by cohesin. Cohesion between sister DNAs was measured by marking the *URA3* locus 35 kb from *CEN4* using a tandem array of Tet operators bound by a GFP-tagged Tet repressor. When released at the restrictive temperature (35.5°C), replication gives rise predominantly to cells with a pair of separated GFP spots, indicating lack of cohesion (Michaelis et al., 1997). In contrast, when released at 23°C, replication gives rise to cells with a pair of *URA3* loci that are so close together that only a single GFP dot is visible. This confirms that Scc2 is indeed essential for establishing cohesion and that *scc2-4* is a *bona fide* temperature sensitive allele. Importantly, release of wild type *SCC2* cells gives rise predominantly to cells with a single GFP dot at both temperatures (results not shown). Due to cleavage of most Scc1 during the preceding anaphase, rather little cohesin is associated with the chromosomes of α-factor arrested cells and loading of Scc1 synthesised de novo following release therefore depends on Scc2.

To address whether cohesin loaded at 23°C is sufficient to build cohesion at the restrictive temperature (35.5°C), we released cells from pheromone into medium containing hydroxyurea (HU) at 23°C, which blocks (or rather greatly delays) DNA replication but nevertheless permits the burst of *de novo* cohesin loading normally occurring during late G1. After 45 min, cells were transferred to medium lacking HU at 23°C (Fig. 1a), which permitted cells to complete DNA replication (Fig. 1b). Despite loading of cohesin during the 45 min incubation at 23°C, *scc2-4* cells largely failed to establish sister chromatid cohesion when transferred to HU-free medium at restrictive temperature (Fig. 1c). Thus, loading of cohesin in late G1 does not permit cells to generate sister chromatid cohesion during S phase - in the absence of Scc2 activity. It is important to point out that this experiment does not distinguish whether this is due to a requirement of Scc2 during S phase itself or whether Wapl-mediated turnover (Chan et al., 2012; Kueng et al., 2006) ensures that Scc2 is continually required to maintain cohesin on chromosomes until the point cells initiate DNA replication.

Because a different conclusion concerning Scc2’s role during DNA replication has been drawn from similar experiments using a longer incubation period in the presence of HU (Lengronne et al., 2006), we repeated our experiment, in this case leaving cells in HU for 90 min at 23°C before releasing cells into HU-free medium at 35.5°C. As previously reported and in contrast to the 45 min HU incubation, this permitted most *scc2-4* cells to establish cohesion at the restrictive temperature (Fig. 1c). It is known that cells arrested for longer (90 min) periods in HU manage to initiate replication from early origins and build substantial replication forks (Feng et al., 2006). To investigate whether this induces acetylation of Smc3 and thereby an association between cohesin and DNA that is refractory to Wapl, we used Western blotting to measure Smc3 acetylation following release of cells from pheromone at 23°C in the presence or absence of HU. This revealed very little acetylation by 45 min but substantial acetylation by 90 min (Fig. 1d), raising the possibility that limited replication accompanied by acetylation of Smc3 by Eco1 and not loading per se is what enables cells to build cohesion without further Scc2 activity. To test this, we repeated our temperature shift experiments with ts *eco1-1* (K15285) and *scc1-73* (K15031) cells (Fig. 1c). *SCC1* is required to maintain cohesion as well as to create it. As a consequence, *scc1-73* cells failed to establish cohesion either when shifted at 45 min or at 90 min. In contrast, the behaviour of *eco1-1* cells resembled that of *scc2-4* cells, namely about half the cells shifted at 90 min established stable cohesion at 35.5°C while few if any did when shifted at 45 min.

**Figuer 1:**
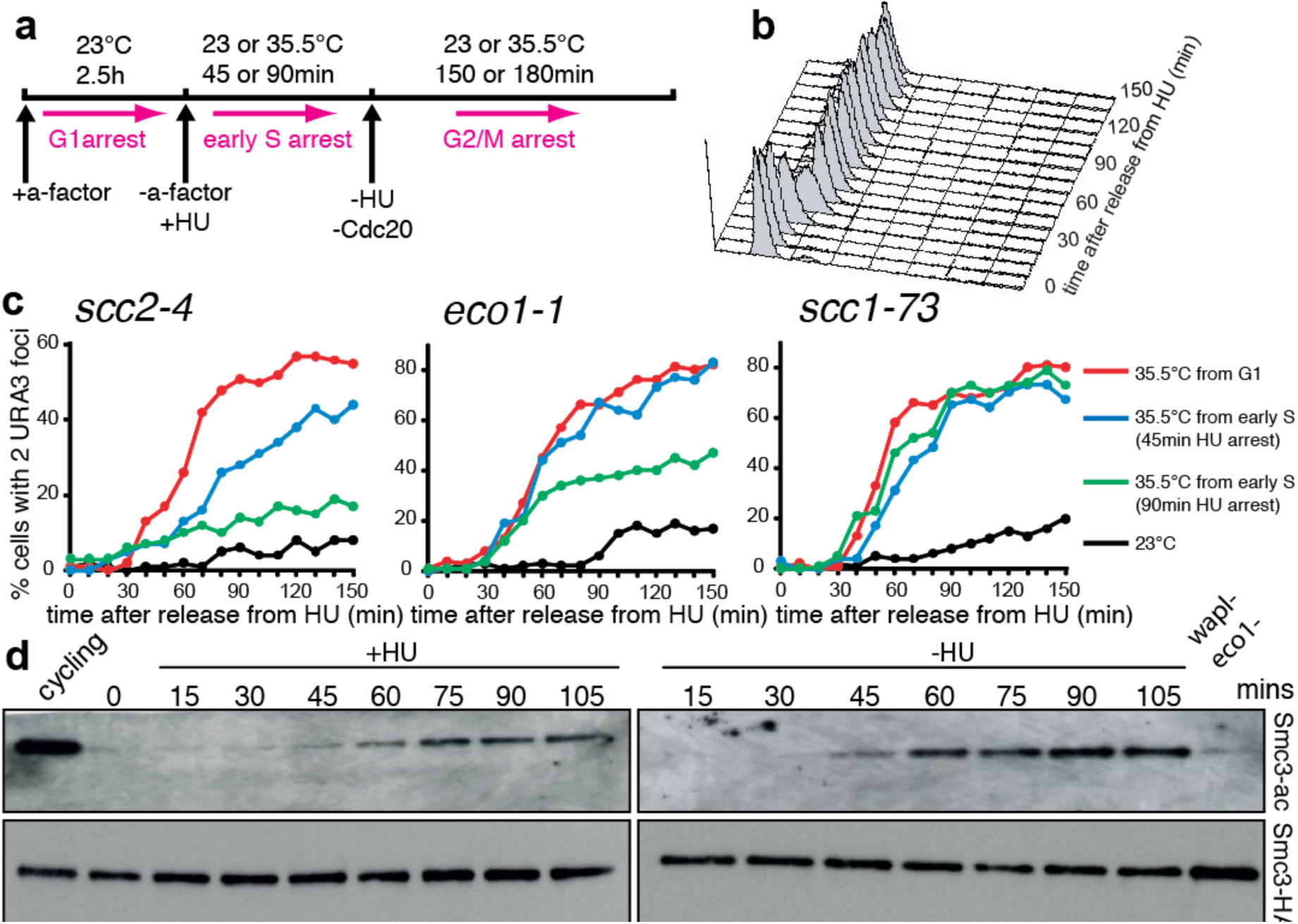
*S.cerevisiae* cells arrested in hydroxyurea (HU) generate sister chromatid cohesion. **a)** Schematic of experimental protocol **b)** FACS analysis of cells released from HU. **c)** Percentage of sister chromatid separation measured by counting the fraction of cells with double (split) GFP dots at the URA3 locus In temperature sensitive scc2-4, eco1-1 or scc1–73 mutants. Cells growing at the permissive temperature (23 °C) were first arrested in G1 by a factor and released at permissive temperature (black), restrictive temperature (red line) or into HU. Cells were held in HU for 45 (blue) or 90 (green) minutes before being moved to the restrictive temperature (35.5°C). All cells were then arrested in prometaphase in nocodazole for microscopy, **d)** Immunoblot of acetylated SMC3-K112/113 of cells arrested for different times in HU

Because acetylation accompanies replication, it is difficult to know whether it is replication or acetylation per se that confers cohesion. Crucially, because substantial replication and presumably acetylation of Smc3 occurs in the vicinity of *URA3* in cells arrested in HU for 90 min, it is impossible to exclude the possibility that cohesion is in fact fully established on replicated parts of the genome and that this cohesion is either sufficiently close to *URA3* to hold sister DNAs together upon release or is capable of translocating into the locus upon release. We suggest that contrary to what has been previously claimed (Lengronne et al., 2006), it is not possible to ascertain from these experiments whether cohesin rings that have loaded onto unreplicated chromatin can create cohesion without further Scc2 activity.

We suggest that future experiments exploring the role of Scc2 during replication would need to address the consequences of inactivating Scc2 as cells enter S phase in cells that had been allowed to load cohesin during late G1. Importantly, such experiments would need to be performed in cells lacking Wapl activity, which would otherwise remove cohesin loaded during G1.

Though our experiments do not permit any definitive conclusion regarding Scc2’s role during S phase, they nevertheless reveal that a previous conclusion to the contrary (Lengronne et al., 2006) must now be regarded as premature.

## Experimental Procedures

The yeast strains used in the split dot assay are derivatives of W303 (K699). The Cells were cultured at 25°C in YEP medium with 2% glucose unless stated otherwise. Hydroxyurea was used at 0.1 M. Strains used were as follows:

| Strain ID | Genotype |
| --- | --- |
| K15021 | MATa<br>scc2-4<br>KanMX::Scc1-PK9<br>HIS3::tetR-GFP<br>URA3::tetOs<br>TRP1::Met3-Cdc20 |
| K15031 | MATa<br>scc1-73<br>LEU2::tetR-GFP<br>URA3::tetOs<br>TRP1::Met3-Cdc20 |
| K15285 | MATa<br>eco1-1<br>LEU2::tetR-GFP<br>URA3::tetOs<br>TRP1::Met3-Cdc20 |

## Acknowledgments

We are grateful to K. Shirahige for supplying anti-acetylated Smc3 antibody. The experiment in Figure 1 a-c was performed by Maria Demidova. The experiment in Figure 1 d was performed by Frederic Beckouët.

